# The P-element strikes again: the recent invasion of natural *Drosophila simulans* populations

**DOI:** 10.1101/013722

**Authors:** Robert Kofler, Tom Hill, Viola Nolte, Andrea Betancourt, Christian Schlötterer

## Abstract

The P-element is one of the best understood eukaryotic transposable elements. It invaded *Drosophila melanogaster* populations within a few decades, but was thought to be absent from close relatives, including *D. simulans*. Five decades after the spread in *D. melanogaster*, we provide evidence that the P-element has also invaded *D. simulans*. P-elements in *D. simulans* appears to have been acquired recently from *D. melanogaster* probably via a single horizontal transfer event. Expression data indicate that the P-element is processed in the germline of *D. simulans*, and genomic data show an enrichment of P-element insertions in putative origins of replication, similar to that seen in *D. melanogaster*. This ongoing spread of the P-element in natural populations provides an unique opportunity to understand the dynamics of transposable element spreads and the associated piRNA defense mechanisms.

## Introduction

The P-element is one of the best understood eukaryotic transposable elements and was originally discovered as the causal factor for a syndrome of abnormal phenotypes in *Drosophila melanogaster*. Crosses in which males derived from newly collected strains were mated with females from long established lab stocks produced offspring with spontaneous male recombination, high rates of sterility and malformed gonads, *i.e*. ”hybrid dysgenesis” (Hiraizumi, 1971; Hiraizumi et al., 1973; Kidwell et al., 1977; Engels and Preston, 1979). Eventually it was discovered that hybrid dysgenesis was due to the presence of a transposable element, the P-element (Bingham et al., 1982; O’Hare and Rubin, 1983), which rapidly became the workhorse of *Drosophila* transgenesis (Rubin and Spradling, 1982; Spradling et al., 1995; Bingham et al., 1982; O’Kane and Gehring, 1987). Surveys of strains collected over 70 years show that the P-element spread rapidly in natural *D. melanogaster* populations, between 1950 and 1990 (Kidwell, 1983; Anxolabéhère et al., 1988; Bonnivard and Higuet, 1999). Surveys of other *Drosophila* species revealed that the P-element had been horizontally transferred (HT) from a distantly related species, *D. willistoni* (Daniels et al., 1990) and that it was absent from close relatives of *D. melanogaster*, including *D. simulans* (Brookfield et al., 1982; Engels, 1992; Vieira et al., 1999; Engels, 1983; Daniels et al., 1990). The failure of the P-element to invade *D. simulans* is surprising as both species are cosmopolitan, mostly sympatric, and share insertions from many TE families *via* horizontal transfer (Sánchez-Gracia et al., 2005; Bartolomé et al., 2009). Furthermore, when artificially injected, the P-element can transpose in *D. simulans*, albeit at a reduced rate (*e.g*. Kimura and Kidwell, 1994).

## Results and Discussion

### The recent invasion of *D. simulans* populations

Here, we show that the P-element has recently invaded natural *D. simulans* populations. We sequenced *D. simulans* collected from South Africa (in 2012) and from Florida (in 2010) as pools [Pool-seq; (Schlötterer et al., 2014)] and analyzed TE insertions in these samples using the method of Kofler et al. (2012). We found P-element insertions at 624 sites in the South African sample, and at nine sites in the population from Florida (fig. 1A), with most insertions segregating at low frequencies (< 0.1; supplementary fig. 3). We compared these results to those from *D. melanogaster* samples collected from South Africa (in 2012) and Portugal (in 2008). In *D. melanogaster* the average number of P-element insertions per haploid genome is similar for the two populations (62 in South Africa, 60 in Portugal). In contrast, the two *D. simulans* populations are very different, with 29 P-element insertions found per haploid genome in the South African sample *vs*. 0.4 in Florida. These differences suggest that we sampled the Florida (2010) population in the early phase of P-element invasion and the South African population (2012) at a more advanced phase. Consistent with a recent invasion, we did not find any P-element insertions in a pool of African (Sub-Saharan) *D. simulans* flies sampled between 2001 and 2009 (Nolte et al., 2012). Using sequences from individual flies, we confirmed that the presence of the P-element in *D. simulans* is not due to a low level of contamination of the pooled flies with *D. melanogaster*, or to a technical artifact. We crossed 12 *D. simulans* Florida males to females of the sequenced *D. simulans* reference strain M252 (Palmieri et al., 2014), which lacks the P-element (Kofler et al., 2014), and sequenced single F1 females from these crossses. Progeny of three of these crosses had P-element insertions (9 in cross 116; 2 in cross 174; 12 in cross 211), a subset of which were validated using PCR (supplementary results 3.1).

**Figure 1.**
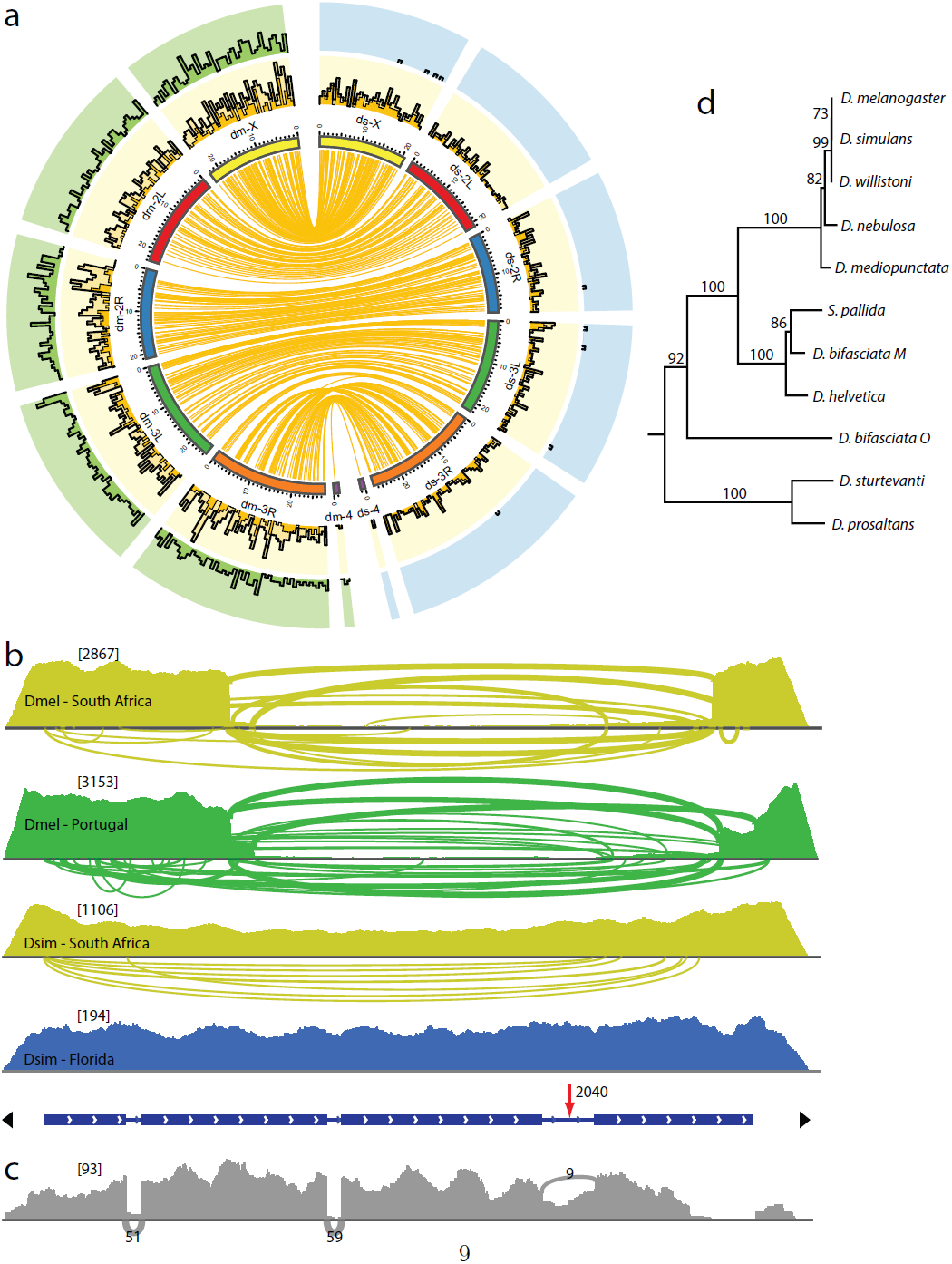
Population genomics of the P-element a) Abundance of P-element insertions in natural *D. simulans* (ds) and *D. melanogaster* (dm) populations. The abundance (in insertions per 500kbp window) is shown for populations from South Africa (yellow), Portugal (green) and Florida (blue). Insertions at similar positions in South Africa are shown in the inlay (dark yellow lines) and the abundance of these insertions is summarized in the histogram (dark yellow). All histograms have a maximum height of 18 insertions. b) Diversity of P-element insertions in natural populations of *D. melanogaster* (Dmel) and *D. simulans* (Dsim). We use Sashimi-plots, which usually indicate splicing in RNA-seq data, to visualize truncated P-element insertions in genomic DNA. Numbers in square brackets are the maximum coverages. The width of the arches scales with the logarithm of the number of reads supporting a given truncated insertion; only truncated insertions supported by at least 3 reads are shown. Panel at the bottom indicates the structure of the P-element (O’Hare and Rubin, 1983). The four ORFs (blue), the SNP distinguishing the *D. simulans* and *D. melanogaster* P-element (red arrow) and the TIRs (black triangles) are shown. c.) Expression of the P-element in *D. simulans*. RNA-seq for the population from Florida was performed and results are visualized with a Sashimi-plot (see above). d.) Phylogeny of the P-element. The P-element of *D. simulans* most closely resembles the P-element from *D. melanogaster* (1 base substitution) and D. willistoni (2 base substitutions).

### Origin of the *D. simulans* P-element

The *D. melanogaster* and *D. simulans* P-element sequences, differ by a single substitution at position 2040 (*G* → *A*; supplementary table 1), which occurs in all *D. simulans* P-element insertions in both populations (supplementary table 1). To identify the origin of the *D. simulans* P-element we constructed a phylogeny using this sequence and that of 10 P-elements most closely related to that of *D. melanogaster* (Clark and Kidwell, 1997, fig. 1d). The *D. simulans* P-element is most similar to the *D. melanogaster* P-element (with 1 nucleotide difference) followed by the *D. willistoni* P-element (2 nucleotide differences), suggesting that *D. melanogaster* is the likely source of the *D. simulans* P-element. If this scenario is true, the *D. simulans* allele at position 2040 might segregate in *D. melanogaster* populations. We screened several publicly available data sets of *D. melanogaster* populations (Kofler et al., 2012, 2014; Bergland et al., 2014) for the presence of this allele, which we found segregating at a low frequency (0.16 – 2%) in 5 of 13 data sets (supplementary results 3.3). This result suggests that the P-element invasion in *D. simulans* was triggered by a single horizontal transfer: recurrent transfer would likely have resulted in the concurrent invasion of the more frequent alternative allele. Consistent with this, P-element insertions in *D. melanogaster* are diverse while those in *D. simulans* are homogenous. *D. melanogaster* P-elements (South Africa π = 0.00072; Portugal π = 0.00121) show a 2 – 3.5 fold higher sequence diversity (Nei and Li, 1979) than P-elements in *D. simulans* (South Africa π = 0.00038; Florida π = 0.00035; see supplementary table 1 for a list of SNPs segregating in P-elements). Further, in *D. melanogaster*, many independently derived truncated P-elements occur [fig. 1b; (O’Hare and Rubin, 1983; Engels, 1992, 1983; Rubin and Spradling, 1982)], while most *D. simulans* P-element insertions are full length (fig. 1b). The predominance of full length insertions in *D. simulans* is consistent with a recent invasion, which requires a functional transposase not encoded by most truncated P-elements (Rubin and Spradling, 1982). In fact, some truncated P-elements repress transposition (Rio, 1990; Black et al., 1987), and may thus inhibit an invasion. The heterogeneity of P-elements in *D. melanogaster* relative to the homogeneity of P-elements in *D. simulans* further supports our hypothesis of a single horizontal transfer event, as recurrent HT probably would have resulted in higher diversity of P-elements in *D. simulans*.

### The *D. simulans* and *D. melanogaster* P-element behave similarly

We analyzed two well-known features of the *D. melanogaster* P-elements, to investigate whether the P-element behaves similarly in *D. simulans*: regulation by alternative splicing and insertion site preferences. In *D. melanogaster* the P-element produces active transposase only in the germline (Engels, 1983), with this tissue specificity controlled post-transcriptionally by alternative splicing of the third intron (Laski et al., 1986). In the soma, transcripts retain the third intron, producing a truncated, inactive version of the transposase protein; in the germ-line, this intron is spliced out yielding a functional transposase (Laski et al., 1986). Host genes responsible for splicing of the third intron (*Psi, Hrb27C*) are highly conserved between *D. melanogaster* and *D. simulans* (Lee and Langley, 2012) and we anticipated that the same pattern of alternative splicing occurs in *D. simulans*. We therefore analyzed RNA-seq data from the Florida *D. simulans* flies for evidence of alternative splicing of the third intron (fig. 1c). We found low levels of spliced transcripts producing transposase, with the splicing of the third intron being supported by 9 reads. However, most reads (38; average of both splice sites) support retention of the third intron, suggesting that the P-element is expressed and regulated somatically in *D. simulans* as in *D. melanogaster* (fig. 1c).

In *D. melanogaster*, the P-element shows a strong preference for insertion into promotor regions of genes (Bellen et al., 2011) and a further bias for origin recognition complex binding sites (Spradling et al., 2011). Here, we found that a substantial fraction of P-element insertions were at similar positions (±1000*bp*) in both *D. melanogaster* and *D. simulans* (428 of 1466 insertions in *D. melanogaster* where 15 are expected due to chance; *χ*^2^ = 11488, *p* < 2.2*e* – 16; fig 1a; supplementary fig. 2). In principle, shared insertions could have been inherited from the ancestor of the two species [2 – 3 million years ago; (Lachaise et al., 1988; Hey and Kliman, 1993)], but this seems unlikely as it would be counter to the evidence showing that the element was absent from both species until recently (Brookfield et al., 1982; Engels, 1992, 1983; Daniels et al., 1990). Further, P-element insertions typically occur at low frequency; it is implausible that the shared insertions have segregated at low frequencies without being lost by genetic drift since the split of the two species. Instead, the presence of insertions at similar sites is likely due to insertion biases: *de novo* P-element insertions tend to occur in a fews hotspots, with 30-40% of all P-element insertions occuring in just 2-3% of the genome (Bellen et al., 2011; Spradling et al., 2011). To investigate whether the same insertion bias occurs in *D. simulans*, we first identified 1kb windows that contained at least 2 independent insertions generated in the course of the Drosophila Gene Disruption Project (18,214 insertions; Bellen et al., 2011). In this way, 2.3% of the genome was identified as potential P-element hotspots (2826 1kb windows). In the *D. melanogaster* sample, 63.5% of P-element insertions from the population from South Africa lie in these regions, representing a significant enrichment (*χ*^2^ = 24226; *p* < 2.2*e* – 16; Table 1). We next identified these hotspots in the *D. simulans* genome using sequence similarity; 54.3% of the *D. simulans* insertions occur at these sites (Table 1), which again is a significant enrichment (*χ*^2^ = 9955; *p* < 2.2*e* – 16). As P-element insertions at similar positions in the two species are significantly more enriched in hotspots (81%) than other P-element insertions (based on *D. melanogaster* insertions; 56.6%; *χ*^2^ = 107; *p* < 2.2*e* – 16), we suggest that the insertion bias accounts for the large fraction of insertions at similar positions. What could be responsible for the insertion bias of the P-element? In fact, the target site specificity of P-elements, which transpose *via* a cut-and-paste mechanism, is thought to be crucial to its invasive properties (Kaufman and Rio, 1992; Engels et al., 1990). That is, its preference for origin recognition complex (ORC) binding sites generates a tendency to insert from replicated into unreplicated DNA, leading to an increase in copy number that does not otherwise occur in cut-and-paste transposition. Consistent with results for *D. melanogaster*, we find the the strongest insertion site bias is for ORCs (Table 1); assuming conservation of ORC sites, we find a 34-fold enrichment of P-element insertions in ORC binding sites (*χ*^2^ = 9703; *p* < 2.2*e* – 16; Table 1).

**Table 1.** Insertion bias of the P-element and other transposable elements in a natural population of *D. melanogaster* (dmel) and *D. simulans* (dsim) from South Africa. Insertions at similar sites in the two species are shown as distinct category (similar). The counts (n) and the relative enrichment relative to a random distribution of insertions in the genome are shown for P-element insertion hotspots, origin recognition complexe binding sites (ORCs) and putative promotor regions (regions within 500bp of a transcription start site). Approximate proportions of genomic features are given brackets. Annotation of features were obtained for *D. melanogaster* and homologous regions in *D. simulans* were identified by sequence similarity. ** highly significant enrichment (*p* < 0.001) relative to other transposable elements

|  |  |  | hotspots (~2%) |  | ORC (~1.6%) |  | promotor (~11%) |  |
| --- | --- | --- | --- | --- | --- | --- | --- | --- |
|  |  |  | n | enrichment | n | enrichment | n | enrichment |
| P-element | dmel | 1,466 | 931 | 27.1** | 672 | 27.3** | 789 | 4.5** |
|  | dsim | 624 | 339 | 31.6** | 319 | 34.3** | 312 | 5.2** |
|  | similar | 428 | 347 | 34.5** | 255 | 35.4** | 234 | 4.6** |
| other transposable elements | dmel | 19,362 | 392 | 0.9 | 290 | 0.9 | 1,791 | 0.8 |
|  | dsim | 13,503 | 143 | 0.6 | 147 | 0.7 | 734 | 0.6 |
|  | similar | 1,017 | 22 | 0.9 | 6 | 0.4 | 116 | 1.0 |

### Reasons for the delayed invasion of *D. simulans*

Why did it take the P-element almost 50 years longer to invade *D. simulans* than to invade *D. melanogaster*? The two hypotheses put forward to explain this phenomenon (Scavarda and Hartl, 1984) invoked either genomic factors that prevent the establishment of the P-element in *D. simulans* (Kimura and Kidwell, 1994; Montchamp-Moreau, 1990) or the rarity of horizontal transfer (Daniels et al., 1985). Genomic barriers to the establishment of P-element in *D. simulans* might have been overcome by adaptation of the TE to *D. simulans*. The single substitution distinguishing the *D. melanogaster* and *D. simulans* P-element seems unlikely to confer a functional advantage: it occurs in an intron, does not coincide with characterized splicing motives (Laski and Rubin, 1989; O’Hare and Rubin, 1983) or with the 9bp motif responsible for maternal transmission (Simmons et al., 2002). Instead, our observations suggest that successful horizontal transmission of the P-element might be rare. The data here suggest that the horizontal transfer to *D. simulans* occured once: recurrent invasion from *D. melanogaster* would result in *D. simulans* insertions harboring a subset of the diversity of the insertions in *D. melanogaster*, or at least the concurrent invasion of the wild-type allele of the *D. melanogaster* P-element. Instead, we find that *D. simulans* insertions are fixed for a rare *D. melanogaster* variant.

## Conclusions

Our observation of a recent, ongoing invasion of transposable elements into a previously uninfected species provides an unique opportunity to study the dynamics of transposable element spread in natural populations. Of particular importance will be the ability to link the ongoing spread with the buildup of piRNAs controlling the spread of P-elements and their relative dynamics in different environments given the strong temperature dependence of hybrid dysgenesis (Engels, 1983). The invasion of *D. simulans* may lead to the P-element rapidly invading the rest of the *melanogaster* subgroup; both P-element free relatives, *D. mauritiana* (where we did not find any P-element insertions; supplementary results 3.4) and *D. sechellia* (Brookfield et al., 1982; Daniels et al., 1990), are known to hybridize with *D. simulans* (Nunes et al., 2010; Matute and Ayroles, 2014).

## Material and Methods

We measured TE abundance in two populations of *D. melanogaster* and two populations of *D. simulans* using Pool-seq data and PoPoolation TE (Kofler et al., 2012) as described in (Kofler et al., 2014). We used three previously published data sets [*D. melanogaster* from South Africa; (Kofler et al., 2014); *D. melanogaster* from Portugal (Kofler et al., 2012); *D. simulans* from South Africa (Kofler et al., 2014)] and additionally sequenced a *D. simulans* population from Florida as pool using Illumina paired-end sequencing. To confirm the presence of the P-element in *D. simulans* we crossed several males from Florida with the *D. simulans* strain M252 (Palmieri et al., 2014) and sequenced F1 progeny individually with Illumina paired-end sequencing. PCR-primers were designed to confirm insertions in the progeny and amplicons were sequenced using the Sanger technology. We used RNA-seq to measure expression of the P-element. RNA was extracted from *D. simulans* females from Florida that were kept in the lab for 2 generations at 15°C. Insertion bias of P-elements was measured using publicly available data of 18,214 independent P-element insertions (Bellen et al., 2011), origin recognition complex binding sites (Spradling et al., 2011), and 500bp within transcription start sites was used as putative promotor sequences (annotation of *D. melanogaster* v5.57; http://flybase.org/). The programming language R (R Core Team, 2014) was used for all statistical analysis. See supplementary data for more details. The Sanger sequenced amplicons and the *D. simulans* P-element can be found at Genbank (http://www.ncbi.nlm.nih.gov/genbank/ amplicons KP241673-KP241675, P-element KP256109). The Illumina reads have been made available at the short read archive (http://www.ncbi.nlm.nih.gov/sra PRJEB7936).

## Acknowledgments

We thank all members of the Institute of Population Genetics for feedback and support. This work was supported by the ERC grant “Archadapt” and Austrian Science Funds (FWF) grant P27048.

